# Computational epitope mapping of class I fusion proteins using Bayes classification

**DOI:** 10.1101/2022.05.23.493016

**Authors:** Marion F.S. Fischer, James E. Crowe, Jens Meiler

## Abstract

Antibody epitope mapping of viral proteins plays a vital role in understanding immune system mechanisms of protection. In the case of class I viral fusion proteins, recent advances in cryo-electron microscopy and protein stabilization techniques have highlighted the importance of cryptic or ‘alternative’ conformations that expose epitopes targeted by potent neutralizing antibodies. Thorough epitope mapping of such metastable conformations is difficult, but is critical for understanding sites of vulnerability in class I fusion proteins that occur as transient conformational states during viral attachment and fusion. We introduce a novel method Accelerated class I fusion protein Epitope Mapping (AxIEM) that accounts for fusion protein flexibility to significantly improve out-of-sample prediction of discontinuous antibody epitopes. Harnessing data from previous experimental epitope mapping efforts of several class I fusion proteins, we demonstrate that accuracy of epitope prediction depends on residue environment and allows for the precise prediction of conformation-dependent antibody target residues. We also show that AxIEM can to identify common epitopes and provide structural insights for the development and rational design of vaccines.

**Author Summary:** Efficient determination of neutralizing epitopes of viral fusion proteins is paramount in the development of antibody-based therapeutics against rapidly evolving or undercharacterized viral pathogens. Advances in the determination of viral fusion proteins in multiple conformations with ‘cryptic epitopes’ during attachment and fusion has highlighted the importance of epitope accessibility due to viral fusion protein flexibility, a physical trait not accounted for in previous B-cell epitope prediction methods. Given the relatively limited number of viral fusion proteins that have been determined in multiple conformations that also have been extensively subjected to epitope mapping techniques,, which are predominantly class I fusion proteins, we chose a limited feature set in combination with a low-complexity Bayesian classifier model to avoid overfitting. We show that this model demonstrates higher accuracy in out-of-sample performance than publicly available epitope prediction methods. Additionally, due to limited structural annotation of neutralizing epitope residues, we provide examples of how our model better discerns conformation-specific epitopes, which is critical for subunit vaccine design, and how this may provide a novel approach to assess the structural changes of antigenicity of viral fusion protein homologues.

## Introduction

Successful structure-based vaccine design relies on the identification of antigenic determinants that are most likely to elicit a humoral immune response, which can be achieved through the process of epitope mapping(1). Given the time and cost of experimental methods used for epitope mapping, computational B-cell epitope prediction may provide a more practical starting point to narrow the search for commonly conserved or novel epitope targets. Although B-cell epitopes are typically defined as a spatially clustered set of residues with a surface area of 600 Å^2^ to 1,000 Å^2^,(2) the precise definition of both an epitope’s size and residue composition are not always readily known. Therefore, the major challenge of B-cell epitope prediction is making a precise and accurate distinction between residues that are or are unlikely to contact an antibody and whether epitope residues are contiguous in sequence or not.

Data collections for structural epitopes, especially residue-specific data, have increased in the past several years through the curation of databases such as the Immune Epitope Data Base (iedb.org). This availability of data has permitted more accurate epitope prediction models, such as those provided by publicly available Discotope or Ellipro discontinuous epitope prediction servers(3,4). Even so, these epitope prediction models are limited to predicting epitopes of a single protein structure or even a single protein chain, which hinders the prediction of quaternary B-cell epitopes. Moreover, proteins are dynamic, *i.e.*, they assume more than one conformation, and a single conformation of a protein may not be sufficient to predict all possible epitopes. For instance, the stabilization of the respiratory syncytial virus (RSV) fusion (F) protein in its meta-stable prefusion conformation coincided with the identification of a broadly-neutralizing epitope designated Site Ø, which is surface-inaccessible in the more stable postfusion F protein conformation(5).

The innate flexibility of class I viral fusion glycoproteins facilitates the entropy-driven process of membrane fusion to achieve cellular entry and host infection despite distinct fusion mechanisms. Compared to other proteins with antigenic determinants within a viral quasispecies, fusion proteins are more frequently targeted by broadly neutralizing antibodies, and therefore are prime candidates for rational structure-based viral vaccine design (so-called ‘reverse vaccinology’). As most viral fusion proteins are oligomeric and flexible, computational B-cell epitope prediction for these targets faces unique challenges. For thorough epitope mapping and prediction, the model should account for not only the prefusion quaternary structure of the target antigen, but also the changes in quaternary structure during attachment and fusion. Recent advances in experimental design and cryogenic electron microscopy (cryo-EM) allow discovery of cryptic epitopes in ‘alternative’ conformations of viral fusion proteins. It is now feasible to identify residue-specific epitope accessibility changes during the fusion process, albeit with great effort for each antibody-antigen interaction.

We developed a machine learning approach designated Accelerated class I fusion protein Epitope Mapping (AxIEM) that harnesses evolutionary and structural features to classify whether a residue will reside within an epitope depending on the conformation of the fusion protein. We applied AxIEM to seven class I viral fusion proteins for which structures have been determined in at least two conformations and have been extensively subjected to experimental epitope mapping techniques. We show that AxIEM enables a much higher out-of-sample success rate in defining viral fusion epitopes than previous methods and provides a computational tool to identify antigenic determinants of novel or under-characterized viral fusion proteins.

## Results

### Description of AxIEM Dataset

The dataset used to build the training and test sets of the AxIEM model included seven class I fusion proteins (**S1 Table**), where each trimeric protein included at least two conformations of greater than 2.00 Å root mean square deviation (RMSD) and no two conformations of less than 1.00 Å RMSD(6–26). For all 46,710 residues within the dataset, each residue possessed an expected classifier label that indicates the residue as an experimentally determined residue to be part of an epitope or not, plus a feature set of four metric values that account for the conformation and associated energetic changes each residue undergoes during various stages of attachment and fusion (**Fig 1**). Three of the four features were calculated to describe the residue alone in terms of its relative surface exposure, stability, and contact changes within the protein ensemble by using the metrics Neighbor Vector (*NV*)(27), the Rosetta-based per-residue total energy score (*REU*) (28), and contact proximity root mean square deviation (*CP_RMSD_*)(29), respectively. Comparison by a Welch’s two-tailed t-test for each residue-specific feature indicated that the mean value differed significantly between residues that have and have not been experimentally determined to form an antibody (Ab) binding interaction, with *p* < 1.00 × 10^-4^ for all three features (**S1 Fig**). Under the assumption that a residue is more likely to form an epitope if its surrounding residues are also likely to do so, we created the Neighbor Sum (*NS*) metric to estimate the antigenicity of the volume surrounding a single residue as a cosine-weighted linear sum of a residue’s own and surrounding *NV, REU, and CP_RMSD_* values, which increased the separation of epitope and non-epitope scores with a Welch’s paired t-test *p* < 1.00 × 10^-10^ (**S2 Fig**). Please refer to Methods and supplementary information for the detailed description of each metric and definition of epitope residues.

**Fig 1.**
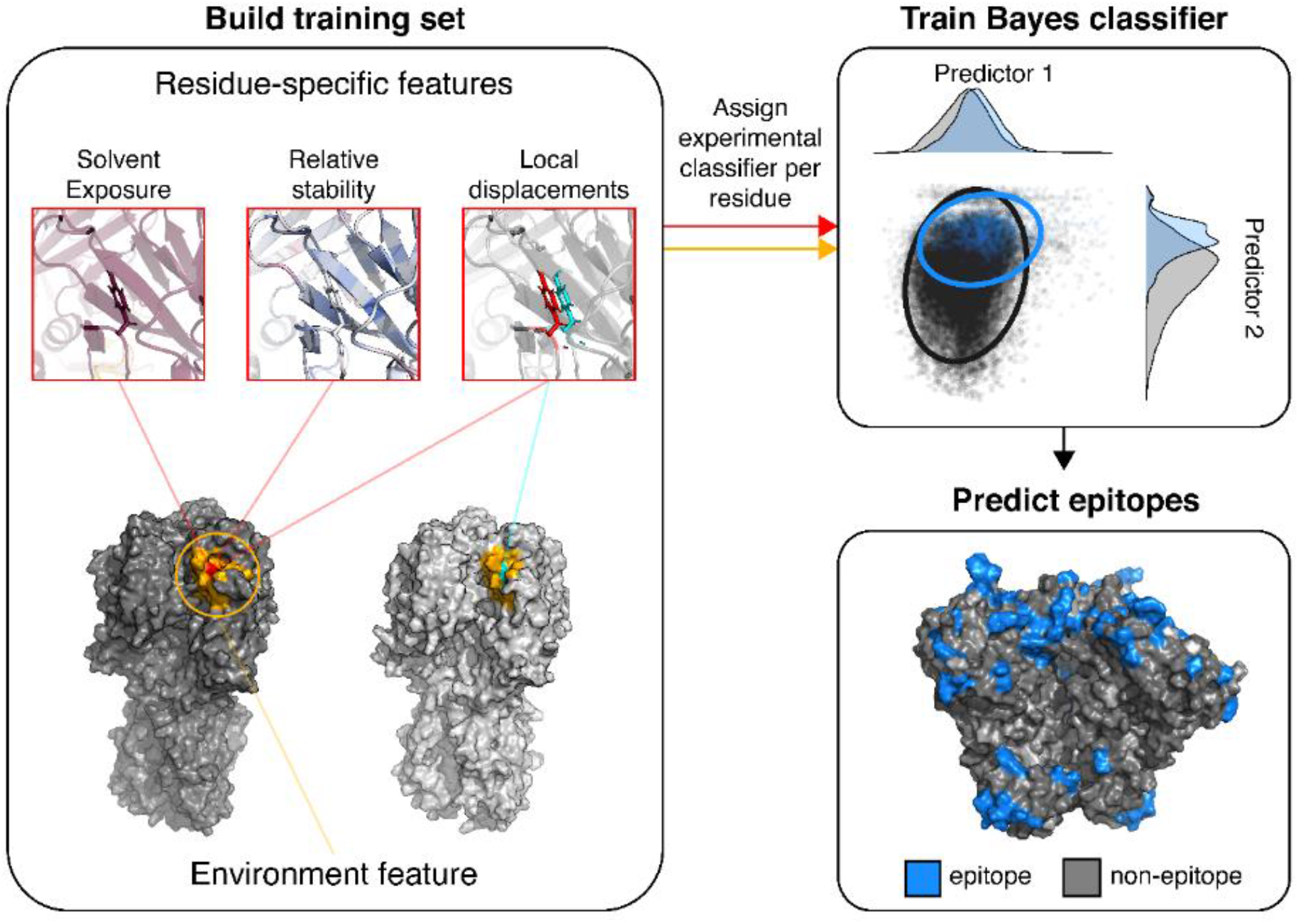
Overview of AxIEM. For each residue within the dataset, a set of four features was calculated that included three residue-specific features (outlined in red) and one environment feature (outlined in dark yellow). Two of the residue specific features that measured relative solvent exposure (as Neighbor Vector) and stability (as Rosetta Energy Unit) are unique to that residue as a part of a single protein conformation. The feature measuring local displacements (as Contact Proximity RMSD) quantifies a residue’s contact changes within an ensemble, and relies on at least two conformations of aligned sequence for calculation. The environment feature (as Neighbor Sum) approximates the relative antigenicity of an area surrounding the residue of interest. A classifier label is assigned to each residue’s calculated feature set for six of the seven protein ensembles. For training the Bayes classifier, the distribution of each of the four features is transformed into a multivariate gaussian distribution, from which a Bayes classifier model is trained to minimize classification error given the training data. For simplicity, only the kernel density estimates of two features are shown. Afterwards, the trained Bayes classifier model is used to predict classification of the left-out protein ensemble. The bottom right panel depicts AxIEM positive predictions for the HIV-1 Env trimer PDB ID 6CM3.

Given the significant differentiation of epitope from non-epitope residues using each metric, we calculated a multivariate Gaussian distribution for each training data’s feature sets and used a Bayes classifier to build a probability model to test for out-of-sample performance on the left-out protein ensemble(30). Performance accuracy relied on the definition of a true positive (*TP*) as a residue with a prediction score above a certain threshold value and also was designated as an expected epitope residue given a specific conformation with a classifier label of ‘1’ prior to building the model. A true negative (*TN*) is a residue with opposite characteristics of a *TP*, although this definition is not as rigid given the possibility of unidentified or incomplete characterization of antigenic sites. A false negative (*FN*) or false positive (*FP*) is any residue that scores incorrectly below or above, respectively, of a given threshold.

### Accuracy of epitope prediction depends on environment score and protein size

We initially chose the metrics that calculated per-residue properties to avoid making assumptions about epitope size or total antigenic surface area. Even though the mean per-residue metric values were significantly different between epitope and non-epitope residues for *NV, REU*, and *CP_RMSD_*, the overlapping coefficient η(31) was high with values of 0.733, 0.685, or 0.792, respectively, indicating that each residue-specific feature has a relatively weak predictive value. With the metric Neighbor Sum (*NS*), we found that increasing the radius size of the neighboring residues’ contributions to the *NS* value of a single residue resulted in a further separation of mean *NS* values between experimentally determined epitope and non-epitope residues up to η = 0.333 at a radius of 40 Å. Evaluation of the receiver operator characteristic (ROC) area under the curve (AUC) values for each viral protein ensemble, however, indicated non-uniform maximal AUC values given the upper boundary radius used to calculate *NS*, despite that cumulative maximal performance converged when using an upper boundary radii greater than 32 Å (**Fig 2**). We found that maximal performance was strongly determined by the number of amino acids within each conformation, with *R^2^ = 0.76* and *p* = 0.011 (**S3 Fig**). Using the linear regression model described by the correlation of protein amino acid number *n* and maximal upper boundary performance *u*, with *u* = 15.1 + 0.0147 *n* to determine the optimal upper boundary radius to calculate *NS* for each protein ensemble, overall performance of the AxIEM method significantly outperformed existing discontinuous epitope prediction methods, Discotope and Ellipro, as summarized in **Table 1**.

**Fig 2.**
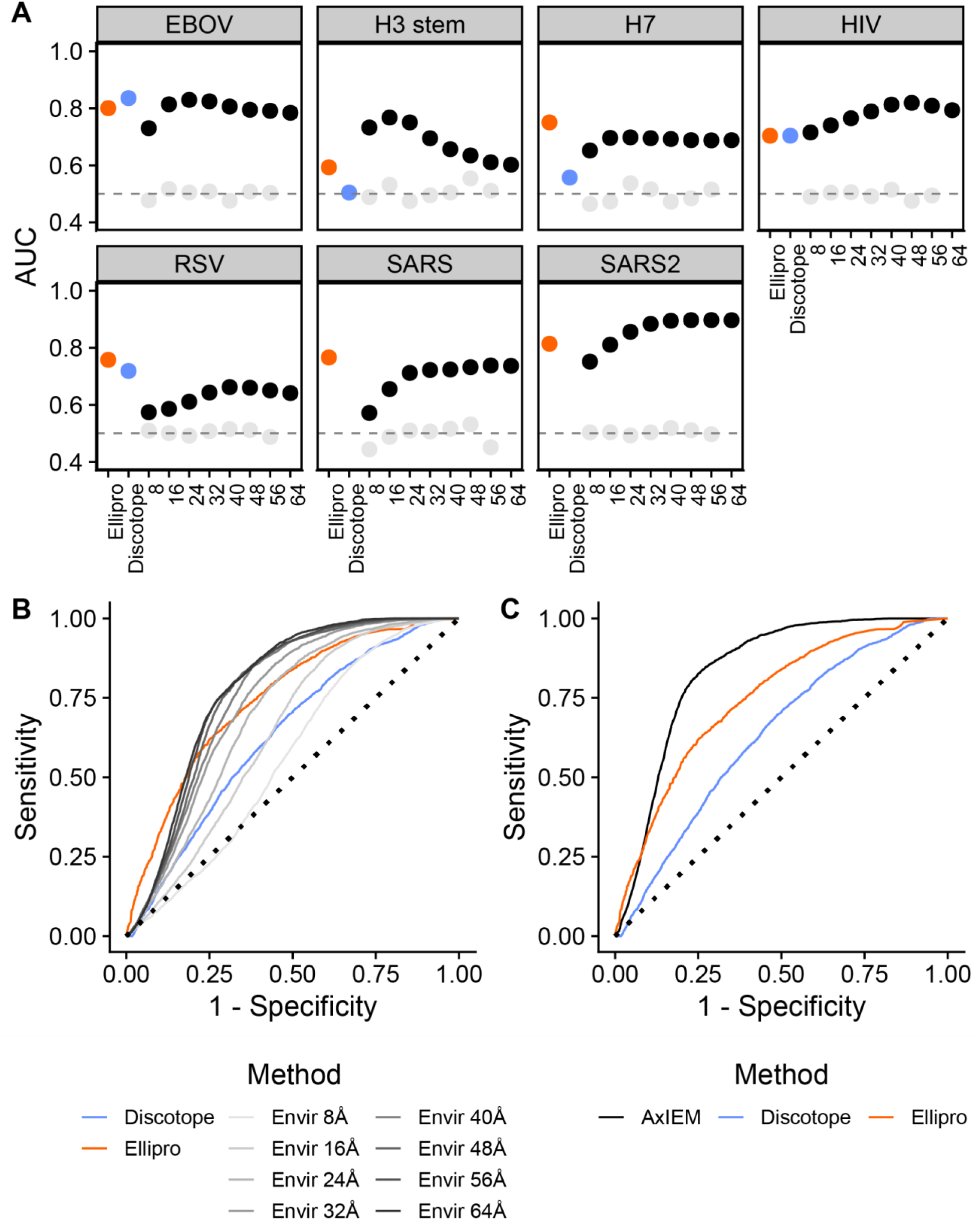
Performance comparison of discontinuous epitope prediction methods. **A)** Comparison of AUC values by virus. Each panel represents the determined AUC value for an individual test set given the employed method. Black indicates that the feature set {*NV, REU, CP_RMSD_, NSu*} was used to train a Bayes classifier model, with *u* as the upper boundary radius listed along the x axis. Light grey represents a negative control for which each residue was assigned a random value for each of the features, *NV, REU*, and *CP_RMSD_* from a normal Gaussian distribution as determined by the mean and standard deviation of each feature’s original values for a single protein. AUC values of Ellipro and Discotope are also listed by protein ensemble test set. Note that Discotope was not able to evaluate predictions for SARS-CoV and SARS-CoV-2 S proteins due to protein size and server time limits. **B)** Comparison of ROC curves by method. Light to dark gray represents increasing upper boundary conditions when used to determine the environment (Envir) feature *NS_u_*. In both panel B and C, the x axis represents the false positive rate calculated as *FP/(FP+TN)*, and the y axis represents the true positive rate calculated as *TP/(TP+FN)*. **C)** Comparison of AxIEM to publically available discontinuous epitope prediction methods. The method AxIEM uses the feature set {*NV, REU, CP_RMSD_, NSu*} where *u* is determined as *u* = 15.1 + 0.0147 *n*, with *n* being the number of residues of a single protein conformation.

**Table 1.**
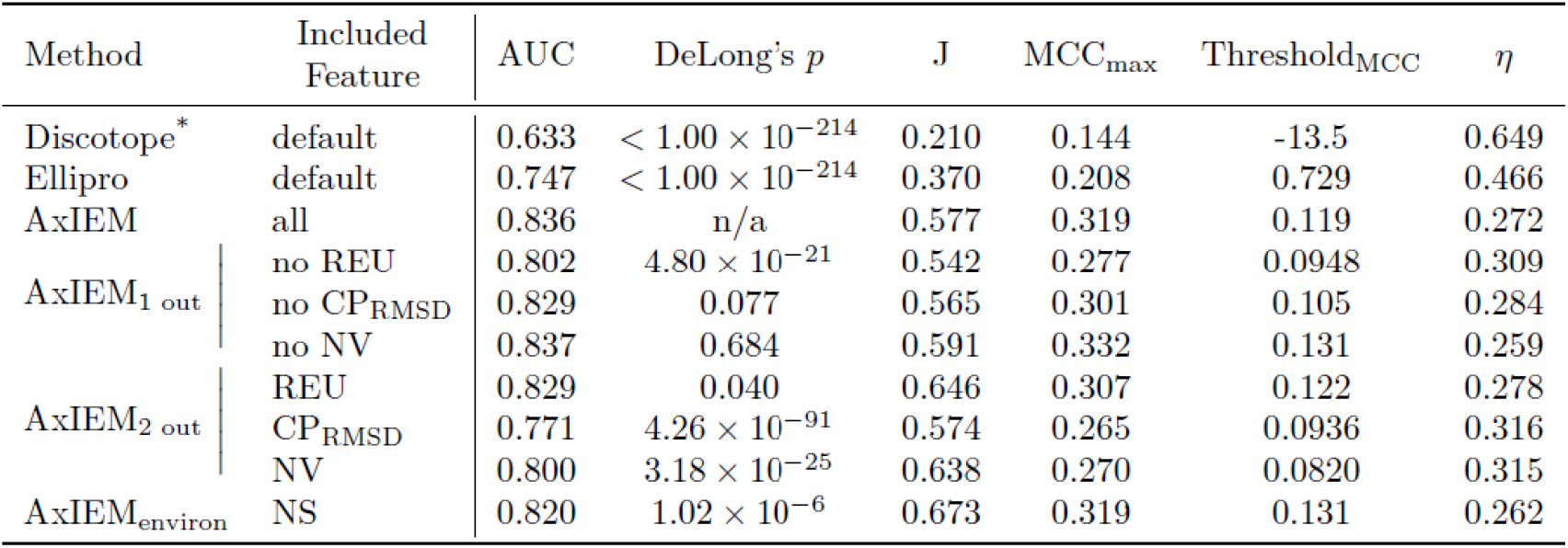
Performance evaluation summary of cumulative predictions. For methods AxIEM_1 out_ and AxIEM_2 out_, one or two of the per-residue features were excluded in the feature set and also from the environment *NS* calculations. The AxIEM_environ_ method excludes all AxIEM per-residue features. Note (*), evaluation of Discotope does not include SARS-CoV S protein epitope predictions.

| Method | Included Feature | AUC | DeLong's $p$ | J | MCC<sub>max</sub> | Threshold<sub>MCC</sub> | $\eta$ |
| --- | --- | --- | --- | --- | --- | --- | --- |
| Discotope* | default | 0.633 | $< 1.00 \times 10^{-214}$ | 0.210 | 0.144 | -13.5 | 0.649 |
| Ellipro | default | 0.747 | $< 1.00 \times 10^{-214}$ | 0.370 | 0.208 | 0.729 | 0.466 |
| AxIEM | all | 0.836 | n/a | 0.577 | 0.319 | 0.119 | 0.272 |
| AxIEM<sub>1 out</sub> | no REU | 0.802 | $4.80 \times 10^{-21}$ | 0.542 | 0.277 | 0.0948 | 0.309 |
|  | no CP<sub>RMSE</sub> | 0.829 | 0.077 | 0.565 | 0.301 | 0.105 | 0.284 |
|  | no NV | 0.837 | 0.684 | 0.591 | 0.332 | 0.131 | 0.259 |
| AxIEM<sub>2 out</sub> | REU | 0.829 | 0.040 | 0.646 | 0.307 | 0.122 | 0.278 |
| | CP<sub>RMSE</sub> | 0.771 | $4.26 \times 10^{-91}$ | 0.574 | 0.265 | 0.0936 | 0.316 |
| | NV | 0.800 | $3.18 \times 10^{-25}$ | 0.638 | 0.270 | 0.0820 | 0.315 |
| AxIEM<sub>environ</sub> | NS | 0.820 | $1.02 \times 10^{-6}$ | 0.673 | 0.319 | 0.131 | 0.262 |

### AxIEM clarifies conformation specificity of common epitopes

Correct assignment of a classifier label, or in other words, what range and combination of feature values the model is instructed to classify as an epitope or not, is crucial in assessing model accuracy. In the case of SARS-CoV or SARS-CoV-2 Spike (S) proteins, most available structures of Ab-antigen interactions exist as fragment Ab and receptor binding domain (RBD) complexes, which disallows the certainty of classifier assignment of experimentally determined epitope residues to any one or more S protein trimeric conformation species. Given the experimental information available and the definition of an epitope residue used for the curation of the AxIEM dataset, the closed conformations of both SARS-related S proteins were labelled to be antigenic, while only the 2- or 3-RBD up or open conformations of the SARS-CoV-2 S protein were labelled to be antigenic. Although the AUC values of S protein-specific predictions were relatively high for SARS-CoV and SARS-CoV 2, with values of 0.759 and 0.908, up to 65.3% of false prediction assessment types could possibly be attributed to incorrect classifier label assignment, suggesting that the feature set used by AxIEM is sufficient to discern relevant conformations that will also give rise to an antigenic response (**Fig 3 and Table 2**). More importantly, AxIEM allows for the comparison of antigenic sites between homologs, since the AxIEM feature set is agnostic to sequence identity. Sequence alignment of predicted positives indicated only eleven aligned predicted positive residues between S protein monomers are structurally available to form an Ab-antigen interaction when in the up or open conformation (**Fig. 3B**). Geographically speaking, aligned ‘up’-conformation positives share a relative constellation pattern, but differ in their relative orientation to each other by a root mean squared difference in pairwise distances of 19.4 Å (as calculated in **Equation 7**. Of the eleven aligned positives, only six residues share the same sequence identify with a root mean squared difference in epitope residues’ pairwise distances of 3.64 Å (**S2 Table**), indicating that RBD epitope similarity is relatively low between SARS-related S proteins. Besides predicting experimentally determined RBD epitopes, AxIEM also predicted a novel common site of vulnerability within the N-terminal domain (NTD) of the S1 subunit for both SARS-CoV-2 and SARS-CoV S proteins, although the sequence identity and alignment differed for all predicted NTD epitope residues (**Fig. 3C**). Given these data, identification of a broadly-neutralizing Ab against multiple SARS-related S proteins is constrained not only by sequence similarity, but also conformation similarity and availability as metastable up or open conformations.

**Fig 3.**
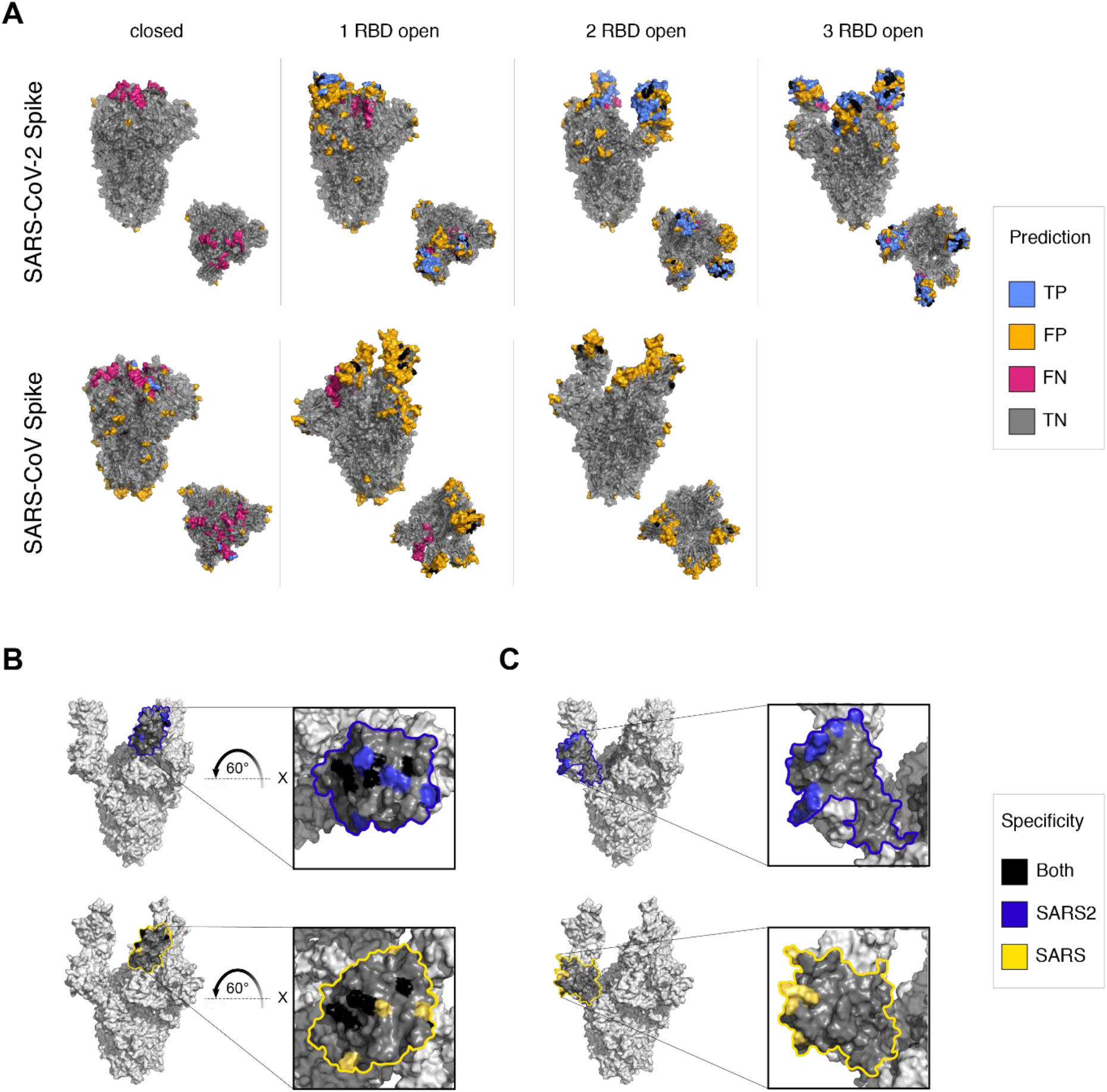
Overview of AxIEM predictions of coronavirus Spike protein epitopes. **A)** Predictions mapped to conformation models. Side and top views are shown for each protein and conformation. Black indicates alignment of positive predictions. **B)** Alignment of common RBD epitopes. Highlighted area represents all residues that are within 16 Å of the geometric centroid of identified common epitopes. The models used to represent the SARS-CoV-2 (top) or SARS-CoV (bottom) include PDB models 7CAK and 6NB7. Black indicates alignment of positive predictions sharing the same sequence identity. Blue and yellow indicate sequence position alignment only. **C)** Alignment of novel NTD epitopes. Highlighted areas, model representation, and coloring are the same as in panel B.

**Table 2.**
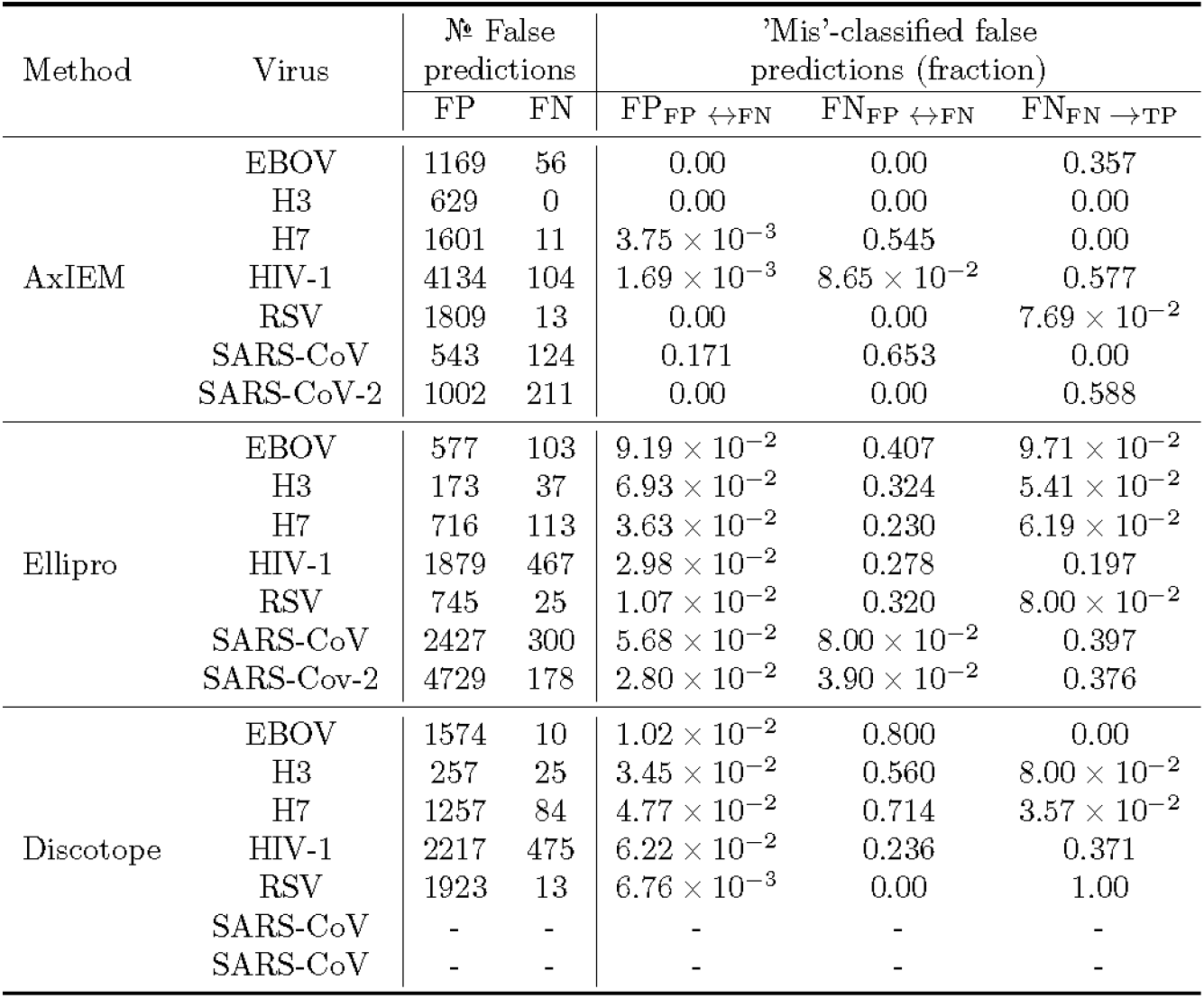
Summary of false predictions. *FP_FP↔FN_* and *FN_FP↔FN_* represents the fraction of *FP* and *FN*, respectively, where for each residue within an ensemble, the residue was labeled as an expected negative for one conformation and predicted to a positive, while also labeled as an expected positive in another conformation that was predicted to be a negaitive. *FN_FN→TP_* represents the fraction of *FN* that were not labeled as expected positives for additional conformations, while in other other conformations that residue was correctly predicted as a positive with no *FP*.

| Method | Virus | № False predictions |  | 'Mis'-classified false predictions (fraction) |  |  |
| --- | --- | --- | --- | --- | --- | --- |
| | | FP | FN | $FP_{FP \leftrightarrow FN}$ | $FN_{FP \leftrightarrow FN}$ | $FN_{FN \rightarrow TP}$ |
| AxIEM | EBOV | 1169 | 56 | 0.00 | 0.00 | 0.357 |
|  | H3 | 629 | 0 | 0.00 | 0.00 | 0.00 |
| | H7 | 1601 | 11 | $3.75 \times 10^{-3}$ | 0.545 | 0.00 |
| | HIV-1 | 4134 | 104 | $1.69 \times 10^{-3}$ | $8.65 \times 10^{-2}$ | 0.577 |
| | RSV | 1809 | 13 | 0.00 | 0.00 | $7.69 \times 10^{-2}$ |
|  | SARS-CoV | 543 | 124 | 0.171 | 0.653 | 0.00 |
|  | SARS-CoV-2 | 1002 | 211 | 0.00 | 0.00 | 0.588 |
| Ellipro | EBOV | 577 | 103 | $9.19 \times 10^{-2}$ | 0.407 | $9.71 \times 10^{-2}$ |
| | H3 | 173 | 37 | $6.93 \times 10^{-2}$ | 0.324 | $5.41 \times 10^{-2}$ |
| | H7 | 716 | 113 | $3.63 \times 10^{-2}$ | 0.230 | $6.19 \times 10^{-2}$ |
| | HIV-1 | 1879 | 467 | $2.98 \times 10^{-2}$ | 0.278 | 0.197 |
| | RSV | 745 | 25 | $1.07 \times 10^{-2}$ | 0.320 | $8.00 \times 10^{-2}$ |
| | SARS-CoV | 2427 | 300 | $5.68 \times 10^{-2}$ | $8.00 \times 10^{-2}$ | 0.397 |
| | SARS-Cov-2 | 4729 | 178 | $2.80 \times 10^{-2}$ | $3.90 \times 10^{-2}$ | 0.376 |
| Discotope | EBOV | 1574 | 10 | $1.02 \times 10^{-2}$ | 0.800 | 0.00 |
| | H3 | 257 | 25 | $3.45 \times 10^{-2}$ | 0.560 | $8.00 \times 10^{-2}$ |
| | H7 | 1257 | 84 | $4.77 \times 10^{-2}$ | 0.714 | $3.57 \times 10^{-2}$ |
| | HIV-1 | 2217 | 475 | $6.22 \times 10^{-2}$ | 0.236 | 0.371 |
| | RSV | 1923 | 13 | $6.76 \times 10^{-3}$ | 0.00 | 1.00 |
|  | SARS-CoV | - | - | - | - | - |
|  | SARS-CoV | - | - | - | - | - |

## Discussion

### AxIEM provides a low complexity solution to epitope mapping

For this study, we chose a final model that employs a Bayes classifier, which has been shown to optimally minimize classification error (32), in conjunction with a limited feature set to avoid overfitting from the experimentally validated dataset. Despite its low complexity, the AxIEM model improves prediction of tertiary and quarternary epitopes of class I viral fusion proteins compared to the IEDB sponsored discontinuous epitope prediction methods, Ellipro and Discotope 2.0. The limited computational requirements of AxIEM, either to use as is or retrain, provides an accessible tool for vaccine development strategies such as screening for novel or cryptic antigenic sites of newly determined class I viral fusion proteins, comparing fusion protein homologues as demonstrated in **Fig 3**, or employing AxIEM within subunit vaccine design platforms. The further use of AxIEM as a computational epitope mapping strategy, however, requires further consideration of aspects of viral structural biology and poses future challenges to generalize epitope prediction, as discussed in the following sections.

### Blocking the moving target requires dynamic precision

To model viral fusion protein flexibility, AxIEM requires at least two conformations to represent major conformation populations during fusion protein-mediated cellular entry and relies on the coarse-grained flexibility metric *CP_RMSD_* to estimate cumulative local residue displaced contacts as previously described. Almost certainly, two conformations are insufficient to fully summarize the biophysical changes of viral entry in terms of representing the major subpopulations, dynamics, and various other gradients like pH when entering the host cell.. Exclusion of *CP_RMSD_* from the AxIEM feature set insignificantly affected overall performance (**Table 1**), and therefore could be excluded when only a single conformation is known. However, AxIEM exceeds in classifying protein antigenic residues when prefusion conformation heterogeneity is more thoroughly represented, such as in the case of ebola glycoprotein, HIV Env, or SARS-CoV-related S proteins that have multiple prefusion conformations included in the datasets. In these cases, the diverging overlap of epitope and non-epitope residue feature distributions is more pronounced, with η < 0.300 for all AxIEM features, compared to the cumulate *NS* feature overlap of η = 0.333.

Conversely, the AxIEM model overestimates the probability of antigenicity of postfusion and membrane-proximal external region (MPER) protein surfaces regardless of conformation, as displayed in **S4-8 Fig,** possibly due to the fact that these regions are not truly surface-accessible to an antibody in a cellular environment. This is supported by the findings summarized in **Table 1**, where exclusion of *NV* had no effect on AxIEM’s AUC with De Long’s *p* = 0.684, and explains the poor individual performance of predicting RSV and influenza H3 stem epitopes due to the large fraction of surface-accessible residues within each represented protein ensemble. However, in the case of HIV Env MPER, antibodies like 10E8 require HIV Env to ‘tilt’ in relation to the lipid bilayer to gain surface accessibility (33), and it is possible that similar regions may present true sites of vulnerability. Further validation would require either identification of novel neutralizing antibodies like 10E8 or better quantification of membrane or protein crowding. Furthermore, any information regarding glycosylation patterns was not included in the AxIEM model due to the lack of high resolution (< 3.0 Å) determination of most glycosylation sites, and therefore any predictions made by AxIEM would need to be supplemented with glycosylation modeling to further assess validity of any initial AxIEM predictions. Overall, AxIEM’s performance and other computational epitope prediction methods would likely benefit from modeling of protein target dynamics and major subpopulation states to better interrogate how protein flexibility affects antigenicity and which major subpopulation states are most likely to illicit a strong neutralizing response.

### Subunit vaccine design may improve by considering protein size

Subunit vaccines rely on the adaptive immune response to produce antibodies against a protein domain or scaffold, which can subsequently elicit a neutralizing antibody response during natural infection. Sites of vulnerability are viral protein surfaces that have been shown to form more than one neutralizing antibody binding interaction and are prime candidates for serving as antigen targets for subunit vaccines. Rather than assuming that proximity features for antigenicity prediction are constrained by the surface area of an antibody binding footprint like Ellipro or Discotope 2.0, AxIEM improves epitope residue classification prediction by employing physicochemical proximity features in relation to the total number of residues within a fusion protein (**Fig 2**). Given that AxIEM positive predictions tend to be clustered within local domains, many of which are known to form epitope-paratope interactions with multiple neutralizing antibodies, AxIEM may be better suited for the task of identifying conformation-specific sites of vulnerability than the task of B-cell epitope prediction. In this case, subunit vaccine design strategies might benefit from the construction of antigens that account for the relative structural antigenicity of a fusion protein rather than the highly specific identification of a potential epitope region..

## Methods

### Explanation of epitope predictors

#### Neighbor Vector (NV)

The per-residue solvent-accessible surface area (SASA) metric *NV* approximates the proximity and spatial orientation of surrounding residues to estimate relative surface exposure of a residue, as previously described(27). In brief, *NV* employs the Contact Proximity (*CP*) and Neighbor Count (*NC*) algorithms to calculate the sum of each surrounding residue’s C_β_-C_β_ distance (*d*) unit vector weighted by its likelihood to make a contact with the reference residue, calculated as *CP* (**Equation 1**), and is normalized by the sum of all possible contacts within the residue’s vicinity, calculated as *NC* (**Equation 2**). In other words, for a highly buried residue, the weighted *d* unit vectors of all C_β_-C_β_ distances will be directed outwards in many directions so that its *NV* score ≈ ≅ 0, whereas a highly exposed residue’s weighted *d* unit vectors will be directed uniformly so that its *NV* score ≅ 1. We use the lower and upper boundaries of 4.00 Å and 12.8 Å, respectively, because 4.00 Å was shown to be the optimal lower boundary to accurately assess per-residue SASA using *NC* and 12.8 Å is the maximum length of a C_β_-C_β_ distance where any atom of one amino acid’s side chain has been shown to make a direct interaction with another amino acid’s side chain (**Equation 3**).

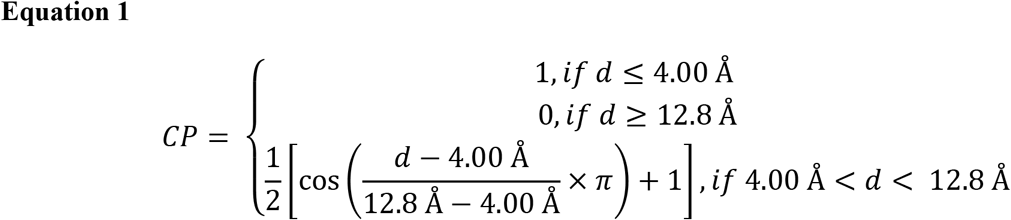

Where *d* is the geometric distance between two C_β_ atoms of two residues. In the case of glycine, a dummy C_β_ atom was used in place of its hydrogen.

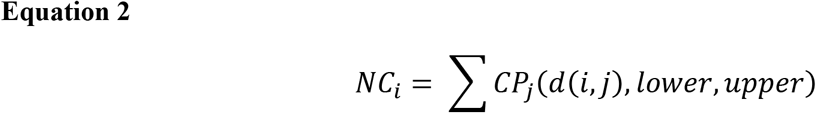

Where *CP_j_* is the evaluated *CP* score of the *j*th residue in relation to the residue of interest *i* given the lower boundary of 4.00 Å and the upper boundary of 12.8 Å.

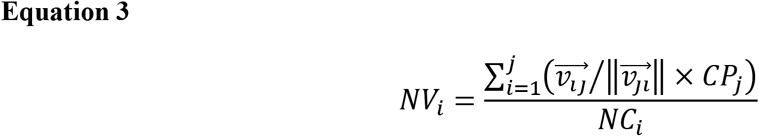

where 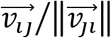 is the unit vector of the *j*th residue multiplied by the *CP* score of the *j*th residue, both terms in relation to the *i*th residue.

#### Per-residue Rosetta Energy Unit (REU)

The relative stability for each residue of a minimized single protein conformation was calculated with the Rosetta *ref2015* energy function and using the jd2_scoring application to estimate the single body and pairwise interaction energies of a residue as the Rosetta per-residue total energy score, which is reported in Rosetta Energy Units (*REU*).

#### Contact Proximity Root Mean Square Deviation (CP_RMSD_)

The metric *CP_RMSD_* has previously been used to estimate the relative local side chain contact changes of a single residue experiences as part of a protein ensemble, and was calculated as the sum of all root mean square deviations of likely contacts a single residue will form (**Equation 4**)(29). This was estimated by *CP* for each C_β_-C_β_ distance, again using 4.00 Å and 12.8 Å as the lower and upper boundaries.

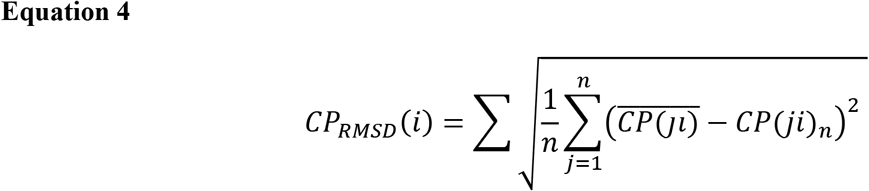

where *i* is the residue of interest, *j* is another residue in the same protein conformation, *n* is the number of protein conformations within the protein ensemble, and 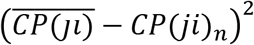 is the variance of the *CP* value of residue *j* with respect to residue *i* within *n* conformations. In the case where only two conformations were used, only the mean *CP* value was calculated.

#### Neighbor Sum (NS)

The *N*S metric was calculated as a weighted linear sum of *NV, REU*, and *CP_RMSD_* (**Equation 5**). In the Results section, we reported *NS* using a fixed lower boundary of 4.00 Å and adjusted the upper boundary to test for epitope radius size as noted (**Equation 6**). We also tested for adjusting the lower boundary to 6.00 Å and 8.00 Å as well as using a linear weighted function instead of a cosine weighted function and found an increase in classification overlap of *NS* values provided that the same upper boundary condition was used.

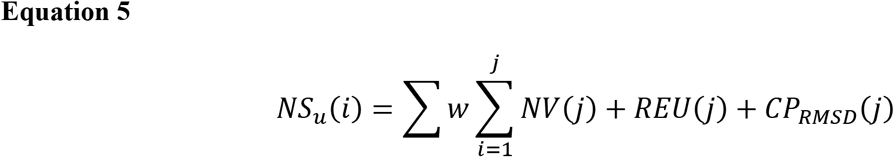

where *i* is the residue of interest, *j* is another residue in the same protein conformation, *u* is the upper boundary radius at which surrounding residues contribute to *i* residue’s *NS* value, and *j* residue’s weighted contribution *w* to residue *i* is determined in **Equation 6**.

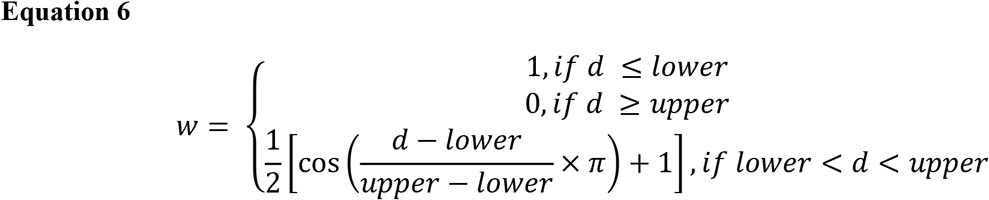

### Definition of conformation-specific epitopes

All residues classified as an “epitope” have been listed within a publicly available online database as experimentally validated discontinuous B-cell epitopes, whose database identifications have been listed in Online Methods. The definition of an epitope was further refined as any residue identified as a discontinuous epitope that also retained a structural similarity to the conformation of the antigen experimentally determined to interact with a specific antibody and that the binding orientation of the antibody did not occupy the same space, or intersect, with any part of the full-length viral protein in that conformation when aligned using PyMOL. Residues that did not fit these criteria were classified as a “non-epitope”.

### Model preparation

Protein structures were downloaded from the Protein Data Bank (PDB; rcsb.org) and aligned using Clustal Omega (www.ebi.ac.uk/Tools/msa/clustalo/) to identify residues that were structurally determined for all equivalent chains and in all conformations of each viral protein. Residues that were not determined as such were manually removed using PyMOL (https://pymol.org/2/) to generate an aligned pdb file. Since some structural models contained mutations for stabilization, the consensus sequence was used to replace the initial sequence. The consensus sequence was obtained by performing a multiple sequence alignment of all available full-length sequences with Clustal Omega and using the EMBOSS cons package (ftp://emboss.open-bio.org/pub/EMBOSS) to identify the consensus sequence. The Rosetta partial_thread application was used to thread, or replace, each residue with the corresponding consensus sequence. Afterwards, the threaded structures were subjected to a constrained relax using the Rosetta FastRelax application to generate 100 models for each protein. From the generated models, the model with the combined lowest energy and lowest structural RMSD to the respective aligned structure was selected as the final model used to evaluate *NV, REU, CP_RMSD_*, and *NS*. For a complete guide to which residues and methods were used for model generation, please refer to the Online Methods section.

### Statistical Information

A DeLong’s test was used to compute significance in differences of AUC between AxIEM performance and other method, where a 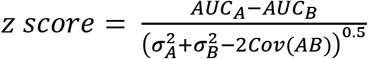. Youden’s J statistic was measured as max (*sensitivity* + *specificity* – 1). A Matthew’s correlation coefficient (*MCC*) was calculated for each threshold along a ROC curve where 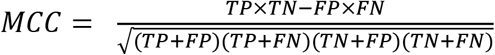. The maximum *MCC* value is reported along with the associated threshold value in **Table 1**. The overlapping index η represents the overlap of prediction scores of expected classifications. Root mean square differences of pairwise epitope geometric distances, denoted as *rmsdp* in **Equation 7**, represents the average difference in geometric distance *d* (**Equation 8**) of each aligned residue *i* to all other residues *j* between epitopes *a* and *b* containing equal number of *n* residues. For a complete description of library and package dependencies and software versions, please see the supplementary protocol capture. All scripts and curated datasets for the development of AxIEM are available at https://github.com/mfsfischer/AxIEM/.

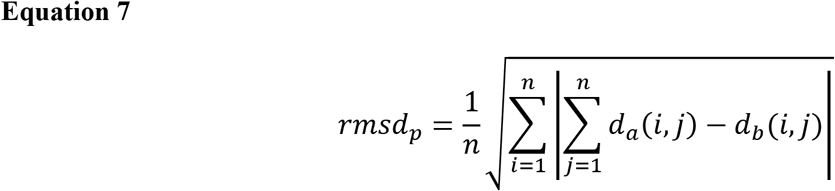

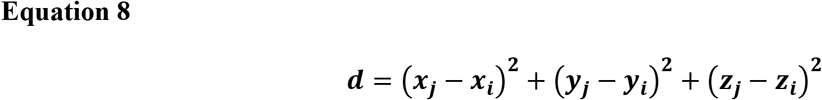

### Computational resources

All calculations were performed on a Core i9-9980HK laptop with 16 GB RAM and Gentoo Linux operating system. Transformation of the feature set to a multivariate Gaussian distribution and training of Bayes classifier models were implemented in Python using the pomegranate package(30). Statistical analysis for ROC curves, AUC values, DeLong’s test(34), Youden’s J index(35), maximal Matthews correlation coefficient(36) and the associated threshold was performed using Python. Conductance of the Welch’s two-tailed t-test, Pearson’s correlation (to determine R^2^, *p* value, and linear model to determine optimal upper boundary radii), and calculation of the distribution-free overlapping index η were performed in R. All datasets, code, and documentation used for this study are publicly available at https://github.com/mfsfischer/AxIEM.

## Acknowledgements

We thank Dr. Axel Fischer for his helpful insights.

## Author contributions

M.F. curated the structure-based epitope data set, implemented code, conducted analysis, and wrote the manuscript. J.M and J.C. contributed to the review of the manuscript.

## Bibliography

1. JE C. Principles of Broad and Potent Antiviral Human Antibodies: Insights for Vaccine Design. Cell Host Microbe. 2017 Aug 9;22(2):193–206.

2. Sela-Culang I, Kunik V, Ofran Y. The structural basis of antibody-antigen recognition. Frontiers in Immunology. 2013;

3. Kringelum JV, Lundegaard C, Lund O, Nielsen M. Reliable B Cell Epitope Predictions: Impacts of Method Development and Improved Benchmarking. Peters B, editor. PLoS Computational Biology. 2012 Dec 27;8(12):e1002829.

4. Ponomarenko J, Bui HH, Li W, Fusseder N, Bourne PE, Sette A, et al. ElliPro: A new structure-based tool for the prediction of antibody epitopes. BMC Bioinformatics. 2008 Dec 2;9(1):514.

5. McLellan JS, Chen M, Chang JS, Yang Y, Kim A, Graham BS, et al. Structure of a major antigenic site on the respiratory syncytial virus fusion glycoprotein in complex with neutralizing antibody 101F. J Virol. 2010;84(23):12236–44.

6. Lee JE, Fusco ML, Hessell AJ, Oswald WB, Burton DR, Saphire EO. Structure of the Ebola virus glycoprotein bound to an antibody from a human survivor. Nature. 2008;

7. Wang H, Shi Y, Song J, Qi J, Lu G, Yan J, et al. Ebola Viral Glycoprotein Bound to Its Endosomal Receptor Niemann-Pick C1. Cell. 2016 Jan;164(1–2):258–68.

8. Bornholdt ZA, Ndungo E, Fusco ML, Bale S, Flyak AI, Crowe JE, et al. Host-Primed Ebola Virus GP Exposes a Hydrophobic NPC1 Receptor-Binding Pocket, Revealing a Target for Broadly Neutralizing Antibodies. Palese P, editor. mBio. 2016 Mar 2;7(1).

9. Bullough PA, Hughson FM, Skehel JJ, Wiley DC. Structure of influenza haemagglutinin at the pH of membrane fusion. Nature. 1994 Sep 1;371(6492):37–43.

10. Weis WI, Brunger AT, Skehel JJ, Wiley DC. Refinement of the influenza virus hemagglutinin by simulated annealing. J Mol Biol. 1990/04/20. 1990;212(4):737–61.

11. Yang H, Chen LM, Carney PJ, Donis RO, Stevens J. Structures of Receptor Complexes of a North American H7N2 Influenza Hemagglutinin with a Loop Deletion in the Receptor Binding Site. Rey FA, editor. PLoS Pathogens. 2010 Sep 2;6(9):e1001081.

12. Turner HL, Pallesen J, Lang S, Bangaru S, Urata S, Li S, et al. Potent anti-influenza H7 human monoclonal antibody induces separation of hemagglutinin receptor-binding head domains. PLoS Biology. 2019;

13. Schommers P, Gruell H, Abernathy ME, Tran MK, Dingens AS, Gristick HB, et al. Restriction of HIV-1 Escape by a Highly Broad and Potent Neutralizing Antibody. Cell. 2020 Feb;180(3):471–489.e22.

14. Gorman J, Chuang GY, Lai YT, Shen CH, Boyington JC, Druz A, et al. Structure of Super-Potent Antibody CAP256-VRC26.25 in Complex with HIV-1 Envelope Reveals a Combined Mode of Trimer-Apex Recognition. Cell Reports. 2020 Apr;31(1):107488.

15. Simonich CA, Doepker L, Ralph D, Williams JA, Dhar A, Yaffe Z, et al. Kappa chain maturation helps drive rapid development of an infant HIV-1 broadly neutralizing antibody lineage. Nature Communications. 2019 Dec 16;10(1):2190.

16. Yang Z, Wang H, Liu AZ, Gristick HB, Bjorkman PJ. Asymmetric opening of HIV-1 Env bound to CD4 and a coreceptor-mimicking antibody. Nature Structural & Molecular Biology. 2019 Dec 2;26(12):1167–75.

17. Wang H, Barnes CO, Yang Z, Nussenzweig MC, Bjorkman PJ. Partially Open HIV-1 Envelope Structures Exhibit Conformational Changes Relevant for Coreceptor Binding and Fusion. Cell Host & Microbe. 2018 Oct;24(4):579–592.e4.

18. Swanson KA, Settembre EC, Shaw CA, Dey AK, Rappuoli R, Mandl CW, et al. Structural basis for immunization with postfusion respiratory syncytial virus fusion F glycoprotein (RSV F) to elicit high neutralizing antibody titers. Proceedings of the National Academy of Sciences. 2011 Jun 7;108(23):9619–24.

19. McLellan JS, Chen M, Joyce MG, Sastry M, Stewart-Jones GB, Yang Y, et al. Structure-based design of a fusion glycoprotein vaccine for respiratory syncytial virus. Science (1979). 2013;342(6158):592–8.

20. Yuan Y, Cao D, Zhang Y, Ma J, Qi J, Wang Q, et al. Cryo-EM structures of MERS-CoV and SARS-CoV spike glycoproteins reveal the dynamic receptor binding domains. Nature Communications. 2017 Apr 10;8(1):1–9.

21. Walls AC, Xiong X, Park YJ, Tortorici MA, Snijder J, Quispe J, et al. Erratum: Unexpected Receptor Functional Mimicry Elucidates Activation of Coronavirus Fusion (Cell (2019) 176(5) (1026–1039.e15), (S0092867418316428), (10.1016/j.cell.2018.12.028)). Vol. 183, Cell. Cell Press; 2020. p. 1732.

22. Walls AC, Park YJ, Tortorici MA, Wall A, McGuire AT, Veesler D. Structure, Function, and Antigenicity of the SARS-CoV-2 Spike Glycoprotein. Cell. 2020 Apr 16;181(2):281–292.e6.

23. Henderson R, Edwards RJ, Mansouri K, Janowska K, Stalls V, Gobeil SMC, et al. Controlling the SARS-CoV-2 spike glycoprotein conformation. Nature Structural & Molecular Biology. 2020 Oct 22;27(10):925–33.

24. Cao Y, Su B, Guo X, Sun W, Deng Y, Bao L, et al. Potent Neutralizing Antibodies against SARS-CoV-2 Identified by High-Throughput Single-Cell Sequencing of Convalescent Patients’ B Cells. Cell. 2020 Jul;182(1):73–84.e16.

25. Chi X, Yan R, Zhang J, Zhang G, Zhang Y, Hao M, et al. A neutralizing human antibody binds to the N-terminal domain of the Spike protein of SARS-CoV-2. Science (1979). 2020 Aug 7;369(6504):650–5.

26. Lv Z, Deng YQ, Ye Q, Cao L, Sun CY, Fan C, et al. Structural basis for neutralization of SARS-CoV-2 and SARS-CoV by a potent therapeutic antibody. Science (1979). 2020 Sep 18;369(6510):1505–9.

27. Durham E, Dorr B, Woetzel N, Staritzbichler R, Meiler J. Solvent accessible surface area approximations for rapid and accurate protein structure prediction. Journal of Molecular Modeling. 2009 Feb 21;15(9):1093–108.

28. Alford RF, Leaver-Fay A, Jeliazkov JR, O’Meara MJ, DiMaio FP, Park H, et al. The Rosetta All-Atom Energy Function for Macromolecular Modeling and Design. Journal of Chemical Theory and Computation. 2017;

29. Sauer MF, Sevy AM, Crowe JE, Meiler J. Multi-state design of flexible proteins predicts sequences optimal for conformational change. PLoS Computational Biology. 2020;

30. Schreiber J. Pomegranate: fast and flexible probabilistic modeling in python. Journal of Machine Learning Research. 2017;18(164):1–6.

31. Pastore M, Calcagnì A. Measuring Distribution Similarities Between Samples: A Distribution-Free Overlapping Index. Frontiers in Psychology. 2019 May 21;10:1089.

32. Theodoridis S, Koutroumbas K. Chapter 2 - Classifiers Based on Bayes Decision Theory. In: Theodoridis S, Koutroumbas K, editors. Pattern Recognition (Fourth Edition) [Internet]. Boston: Academic Press; 2009. p. 13–89. Available from: https://www.sciencedirect.com/science/article/pii/B9781597492720500049

33. Rantalainen K, Berndsen ZT, Antanasijevic A, Schiffner T, Zhang X, Lee WH, et al. HIV-1 Envelope and MPER Antibody Structures in Lipid Assemblies. Cell Rep [Internet]. 2020 Apr 28;31(4):107583. Available from: https://pubmed.ncbi.nlm.nih.gov/32348769

34. DeLong E R, DeLong D M, Clarke-Pearson D L. Comparing the areas under two or more correlated receiver operating characteristic curves: a nonparametric approach - PubMed. Biometrics. 1988;44(3):837–45.

35. Youden WJ. Index for rating diagnostic tests. Cancer. 1950;3:32–5.

36. Matthews BW. Comparison of the predicted and observed secondary structure of T4 phage lysozyme. BBA - Protein Structure. 1975 Oct 20;405(2):442–51.

